# Distributed coding of evidence accumulation across the mouse brain using microcircuits with a diversity of timescales

**DOI:** 10.1101/2022.09.17.506014

**Authors:** Elaheh Imani, Setayesh Radkani, Alireza Hashemi, Ahad Harati, Hamidreza Pourreza, Morteza Moazami Goudarzi

## Abstract

The gradual accumulation of noisy evidence for or against options is the main step in the perceptual decision-making process. Using brain-wide electrophysiological recording in mice (Steinmetz et al., 2019), we examined neural correlates of evidence accumulation across brain areas. We demonstrated that the neurons with Drift-Diffusion-Model-like firing rate activity (i.e., evidence-sensitive ramping firing rate) were distributed across the brain. Exploring the underlying neural mechanism of evidence accumulation for the DDM-like neurons revealed different accumulation mechanisms (i.e. single and race) both within and across the brain areas. Our findings support the hypothesis that evidence accumulation is happening through multiple integration mechanisms in the brain. We further explored the timescale of the integration process in the single and race accumulator models. The results demonstrated that the accumulator microcircuits within each brain area had distinct properties in terms of their integration timescale, which were organized hierarchically across the brain. These findings support the existence of evidence accumulation over multiple timescales. Besides the variability of integration timescale across the brain, a heterogeneity of timescales was observed within each brain area as well. We demonstrated that this variability reflected the diversity of microcircuit parameters, such that accumulators with longer integration timescales had higher recurrent excitation strength.

## 1 Introduction

Decision-making, the process of choosing between options, is a fundamental cognitive function. Different types of decision-making, including perceptual (Gold & Shadlen, 2007) and value-based decision-making (Hunt et al., 2012), is thought to be characterized by a gradual accumulation of noisy evidence for or against options until a threshold is reached and a decision is made. The study of the evidence accumulation process started within cognitive psychology, where researchers explored sequential sampling models, i.e., the drift-diffusion model (DDM), using behavioral data (Ratcliff & McKoon, 2008). In these models, noisy information is accumulated over time from a starting point toward a decision boundary.

Later studies on the neural basis of decision-making developed computational models for the accumulation process using neurons showing signatures of the drift-diffusion model, referred to as DDM-like neurons (Mazurek et al., 2003; Wang, 2002). DDM-like neurons exhibit ramping-like firing rate activity modulated with stimulus coherency. These studies explored some brain regions containing DDM-like neurons, such as the posterior parietal cortex (PPC) (Roitman & Shadlen, 2002; Shadlen & Newsome, 2001), frontal eye field (FEF) (Ding & Gold, 2012; Kim & Shadlen, 1999), striatum (Ding & Gold, 2010), and superior colliculus (Horwitz & Newsome, 1999) in monkeys, as well as the frontal orienting field (FOF) and PPC (Hanks et al., 2015) in rats.

Although previous studies on the neural basis of decision-making explored a few brain regions showing the neural correlate of evidence accumulation, the distribution of DDM-like neurons across the brain is still unknown. Recent brain-wide electrophysiological and calcium imaging studies in mice revealed that neurons involved in decision-making are distributed across the brain (Steinmetz et al., 2019; Zatka-Haas et al., 2021). These findings motivated us to explore the existence of choice-selective neurons that have DDM-like firing rate activity across the brain. Similar to the drift-diffusion process, these neurons have ramping-like firing rates associated with the strength of stimulus evidence, such that stronger evidence levels lead to a faster ramping of firing rate and vice versa. However, these patterns of activity can be explained by different accumulation mechanisms, i.e., single (DDM) and dual accumulators (Bogacz et al., 2006). Although several accumulation models have been proposed in previous studies (Machens et al., 2005; Mazurek et al., 2003; Usher & McClelland, 2001; Wong & Wang, 2006), we examined the popular accumulator circuits (i.e., single and race accumulators) to characterize the underlying neural mechanism of evidence accumulation.

Moreover, the distributed coding of evidence accumulation suggests multiple timescales over this cognitive process (Chen et al., 2015). This property stems from the fact that each brain area exhibits a distinct timescale leading to a hierarchical organization that largely follows the anatomical hierarchy (Chen et al., 2015; Honey et al., 2012; Imani et al., 2023; Murray et al., 2014; Pinto et al., 2022; Rossi-Pool et al., 2021).

As such, we used the brain-wide electrophysiological recording data recently published by (Steinmetz et al., 2019) to investigate the distribution of DDM-like neurons and the underlying neural mechanisms of evidence accumulation across the brain. We demonstrated that evidence accumulation is a distributed process across the brain that is happening through multiple accumulation mechanisms. Our findings revealed that some areas are unilateral and strongly prefer the single accumulation mechanism. On the other hand, some areas are bilateral and contain subpopulations with both single and dual accumulation mechanisms. We further studied the timescale of integration using the simulated data from accumulator models across the brain. The results demonstrated that the accumulator microcircuits have distinct timescales, which were organized hierarchically across the brain, suggesting the existence of evidence accumulation over multiple timescales. Moreover, we observed a heterogeneity of integration timescales within each brain region, reflecting the diversity of recurrent connection strength of the accumulators. Our findings support the hypothesis that microcircuits with longer integration timescales have higher recurrent connection strength.

## 2 Results

### Distributed evidence accumulation across the mice’s brain

To investigate whether or not the evidence accumulation process is distributed across the brain, we used the brain-wide neural recording in mice during a visual discrimination task (Steinmetz et al., 2019). In each trial, a visual stimulus of varying contrast (Gabor patch with sigma 9 and 45° direction) appeared on the right, left, both, or neither side screens. To get a reward, the mice had to turn the wheel to move the stimulus with the higher contrast into the center screen (Figure 1a). During the visual discrimination task, the neural activity of approximately 30,000 neurons in 42 brain areas was recorded using Neuropixel probes. We focused our analysis on the seven groups of brain areas demonstrated in Table 1 and Figure 1b, according to the Allen Common Coordinate Framework (CCF) (Wang et al., 2020).

**Figure 1.**
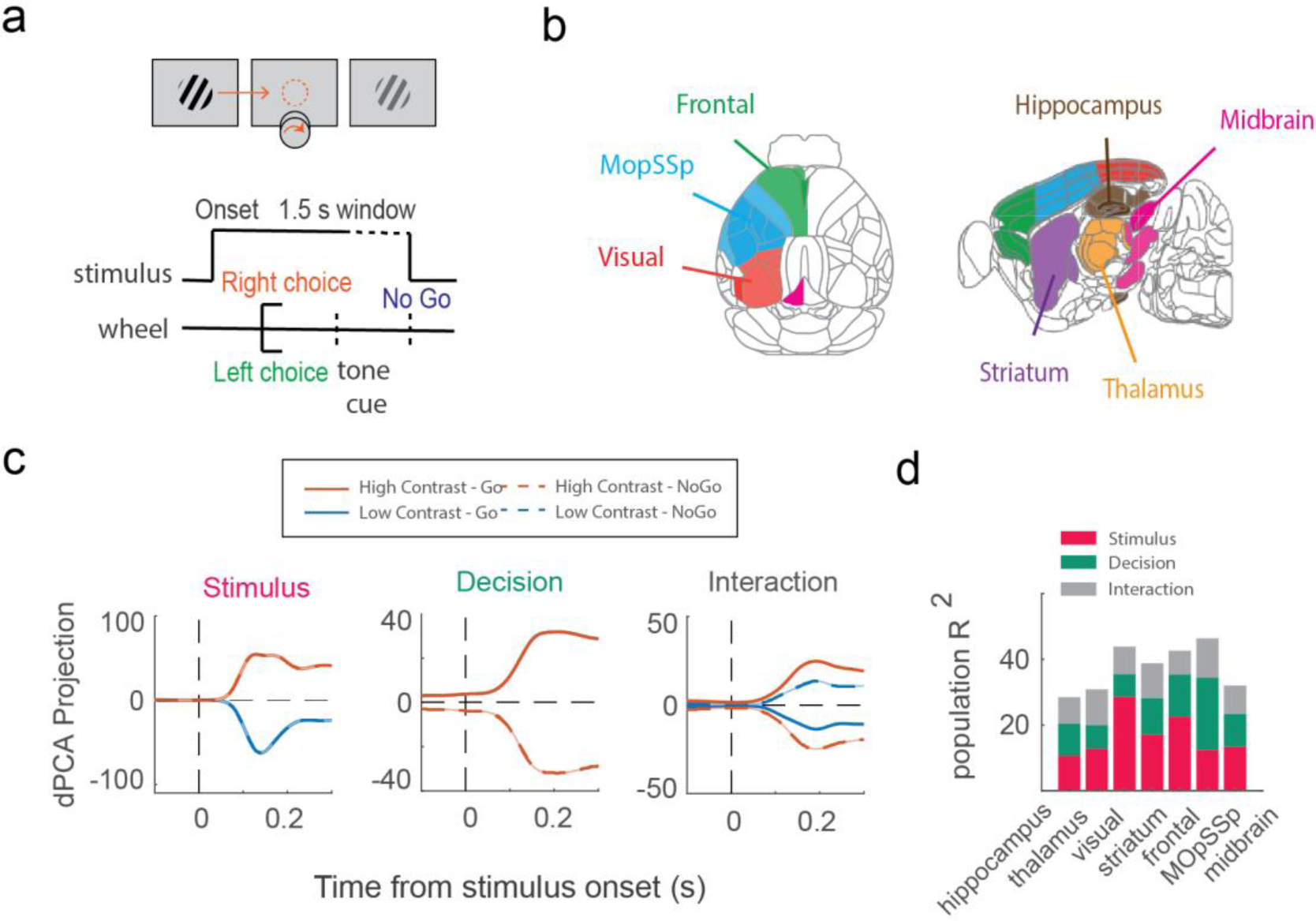
Decomposing brain-wide electrophysiological data into the task-related components using dPCA. (a) Task protocol, (b) Grouping the brain-wide electrophysiological data into the seven regions according to the Allen CCF adapted from (Steinmetz et al., 2019). (c) Projecting the population firing rate into the stimulus, decision, and interaction components. (d) R-squared values of the reconstructed population neural activity using the task-related dPCA components.

**Table 1.** Brain regions within each group of areas according to the Allen CCF.

| Group name | Regions within each group |
| --- | --- |
| Hippocampus | POST, SUB, DG, CA1, CA3 |
| Thalamus | LP, LD, RT, MD, MG, LGd, VPM, VPL, PO, POL |
| Visual | VISp, VISrl, VISam, VISpm, VISl, VISa |
| Striatum | CP, GPe, ACB, LS |
| Frontal | MOs, ACA, PL, ILA, ORB |
| MOpSSp | MOp, SSp |
| Midbrain | MRN, SCm, SCs, APN, PAG, SNr |

To detect the neurons with DDM-like firing rate activity, we first determined the choice-selective neurons within each group of regions. Preliminary analyses showed that most neurons simultaneously encode different task variables, especially in higher cortical areas. Therefore, we first used demixed principal component analysis (dPCA) (Kobak et al., 2016) to decompose the population neural activity into a few principal components representing specific task variables (Figure 1c). We then determined whether a neuron responded more strongly to the stimulus or decision by measuring the reconstructed neural activity’s explained variance (R-squared) using each set of stimulus and decision-related components (supp-Figure 1a). The results revealed that neurons across the brain regions belong to one of three clusters: those best represented by the stimulus-related components, the decision-related components, or their interaction components (Figure 1d, supp-Figure 1b). We excluded the hippocampus region from further analyses because of poor performance in the clustering analysis.

We evaluated dPCA results using the standard auROC metric to measure how well a neuron encodes the stimulus or decision variables. This metric is commonly used to calculate the differences between spike count distributions across different conditions (Britten et al., 1996). There is a strong correlation between the stimulus and animal choice by design. So, we used the combined condition auROC metric to reduce the effect of other task variables on the decoding performance (see Methods). For the stimulus decoding, we measured the differences between the spike count distribution of trials with contralateral stimulus higher than zero and trials with zero contra stimulus contrast level for all 12 conditions.

Similarly, decision decoding was evaluated by measuring the differences between Hit and Missed trials within 12 conditions referred to as ‘Detect Probability’ (DP) (Hashemi et al., 2018). Our results showed that the stimulus-selective neurons detected by dPCA, indeed encoded the stimulus more strongly than the decision. Similarly, the decision-selective neurons encoded the decision better than the stimulus (supp-Figure 1c).

Finally, we found the DDM-like neurons within the decision-related clusters across the brain. Previous studies on the neural basis of evidence accumulation have discovered that DDM-like neurons in the posterior parietal cortex (LIP area) had a ramping-like firing rate activity associated with the strength of a motion stimulus (Roitman & Shadlen, 2002; Shadlen & Newsome, 2001). Similar properties were also found in the mouse’s PPC (Hanks et al., 2015) and anterior dorsal striatum (ADS) in rats (Yartsev et al., 2018). According to the properties of DDM-like neurons, we found the choice-selective neurons that additionally encoded the strength of the input evidence (difference between Right and Left stimulus contrasts). We used the combined condition auROC metric to measure each neuron’s choice probability (CP) and evidence selectivity. Accordingly, we calculated the differences between trials with right and left choices within 12 groups to measure the CP. For measuring evidence selectivity, we evaluated whether or not the trials within a group with the higher evidence level had greater neural activity than those within all the groups having lower evidence (Figure 2b) (See Methods).

**Figure 2.**
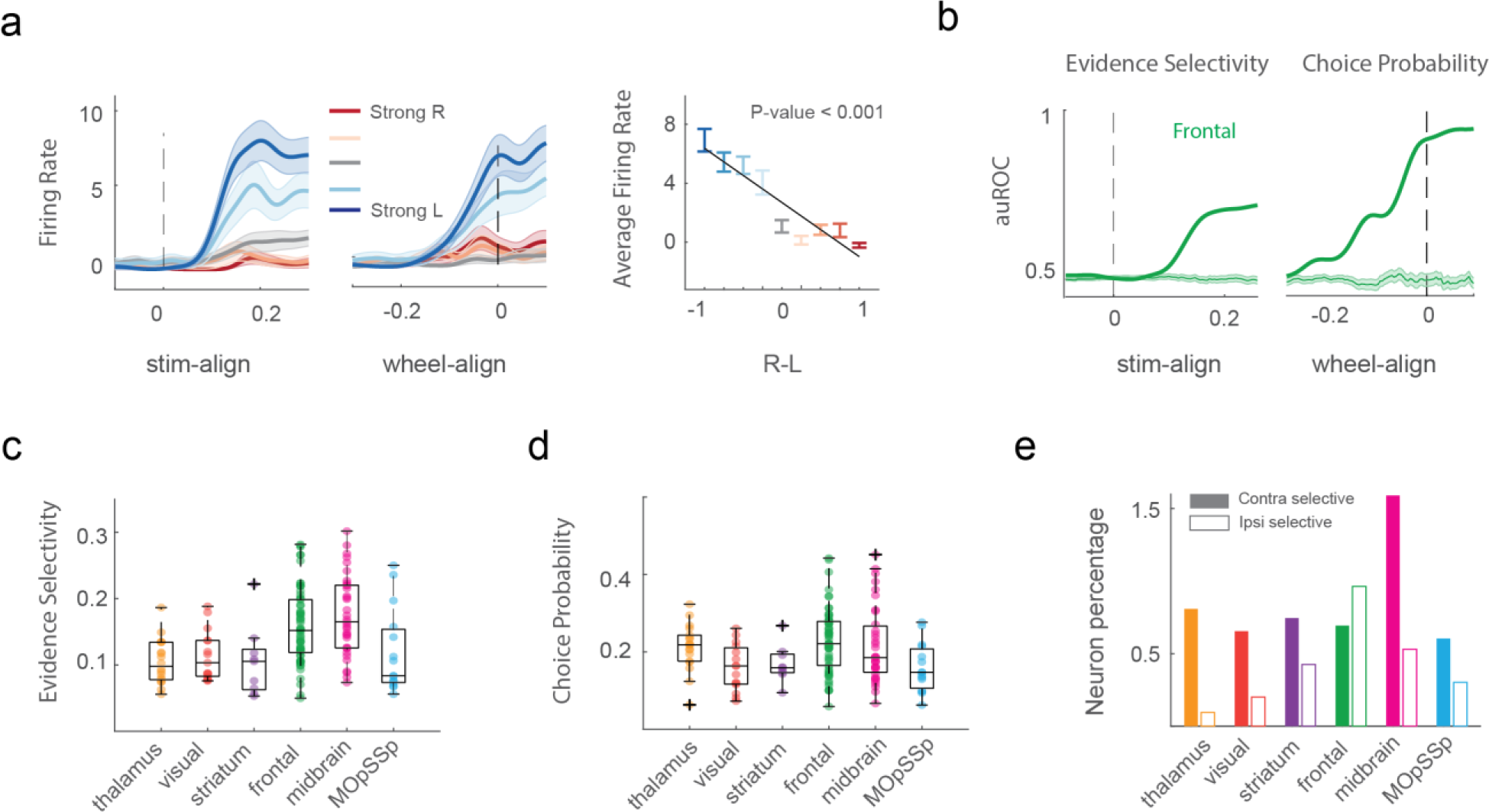
DDM-like neurons across the mouse brain. (a) Example neuron with DDM-like firing rate activity. The curves in the left panel represent neurons’ average firing rate activity across correct trials with a specific evidence level. The color strength indicates the strength of the evidence level, ranging from strong leftward to strong rightward. Shaded areas represent the 95% confidence interval. The right panel shows the linear relationship between the average firing rate of the neuron and the evidence levels using a general linear model. The colors represent the strength of the evidence level and the error bars indicate the 95% confidence interval (b) Temporal evidence selectivity and choice probability for a sample neuron in the frontal region. Panels (c) and (d) are the maximum values of evidence selectivity and choice probability of the DDM-like neurons, respectively. (e) The total number of DDM-like neurons within each brain region. Filled and empty bars represent the number of neurons with contralateral and ipsilateral choice preferences, respectively.

Moreover, to determine whether or not a neuron significantly encoded choice and evidence, we measured decoding performance at the chance level by randomizing the trial labels (Figure 2b). The selective neurons (Figure 2c and 2d) were further visually inspected to exclude those without a ramping-like firing rate activity. The results revealed that the surviving selective neurons have DDM-like firing rate activity (Figure 2a, supp-Figure 2a) and are distributed across the brain regions (Figure 2e, supp-Figure 2b and 2c). Most DDM-like neurons were found in the frontal (MOs, PL, ACA, ILA, and ORB) and midbrain (MRN, SNr, SCm, and SCs) regions. A lower percentage of these neurons were located in the striatum (CP and ACB) and visual pathway (VISam, VISI, and VISp), thalamus (VPL, VPM, LP, PO, LD), and MOpSSp (supp-Figure 2b and 2c). Some of the discovered DDM-like sub-areas within the frontal, striatum, and visual regions were consistent with the previous studies on the neural basis of evidence accumulation in rodents (Hanks et al., 2015; Scott et al., 2017; Yartsev et al., 2018). A single hemisphere contained neurons with both ipsilateral and contralateral choice preferences in most grouped regions (Figure 2e), consistent with the previous studies (Scott et al., 2017). The frontal region was mostly bilateral since the number of the ipsilateral and contralateral DDM-like neurons was similar. In contrast, other brain regions were mostly unilateral.

### Multiple accumulation mechanisms across the brain

Previous studies on the evidence accumulation process proposed different network architectures for evidence integration including single and dual accumulators (Bogacz et al., 2006). Single accumulators such as the drift-diffusion model (DDM) (Ratcliff, 1978) and the ramping model (Latimer et al., 2015; Zoltowski et al., 2019) contain one decision variable accumulating the relative evidence (difference between the two input streams) toward one of the decision boundaries. Dual accumulators are other accumulation mechanisms with separate accumulators for each choice option that integrate the input streams independently (Ditterich et al., 2003; Mazurek et al., 2003) or with mutual inhibitory connections (Machens et al., 2005; Usher & McClelland, 2001; Wang, 2002; Wong et al., 2007; Wong & Wang, 2006). In these accumulation mechanisms, an option is chosen when the integrator associated with that option reaches the decision boundary sooner than the others (Bogacz et al., 2006).

To investigate whether the DDM-like neurons across the mouse brain integrate evidence through a single or dual accumulation mechanism, we used a general framework for the evidence accumulation modeling based on the recurrent switching linear dynamical system (rSLDS) (Zoltowski et al., 2020) (Figure 3). Using rSLDS, the high-dimensional population neural activity can be described as the dynamics of a few continuous latent variables in a low-dimensional state space, evolving through time according to state-dependent dynamic models (Figure 3a). The rSLDS was reformulated to implement the single, independent race, and dependent race accumulation mechanisms (Figure 3b) by considering the accumulators as the continuous latent variables of the model (Figure 3c) (Zoltowski et al., 2020).

**Figure 3.**
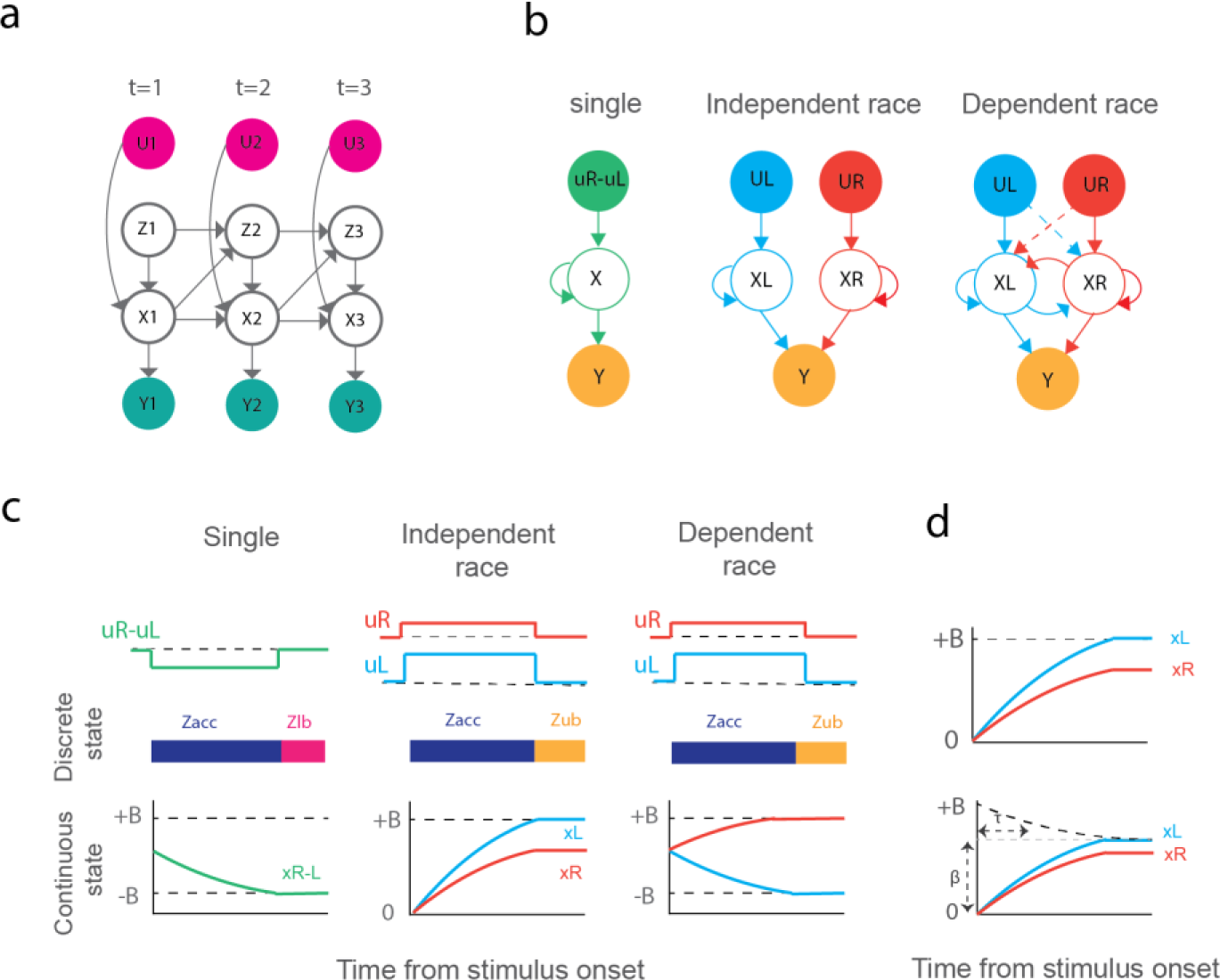
Reformulating the recurrent switching linear dynamical system (rSLDS) framework to the single and dual (independent/dependent race) accumulation mechanisms. (a) Schematic of the rSLDS containing the hidden discrete variable Z, hidden continuous variable X, and observed variables U related to the stimulus strength and neuron spike data Y. (b) Single, independent race, and dependent race accumulator models implemented in rSLDS. (c) Discrete and continuous states of the accumulation mechanisms within sample trials. (d) Constant (top row) and collapsing (bottom row) decision boundaries. The collapsing boundary contains two parameters *β* and *τ*, for the boundary offset and the rate of exponential decay.

We first generated subpopulations of neural activity by resampling the neurons within each region (See Methods). Several bilateral (including neurons with contralateral and ipsilateral choice preference) and unilateral (including neurons with contralateral choice preference) subpopulations were generated during the resampling process (Figure 4c) (See Method). We fit the single and race accumulators to the bilateral subpopulations since these subpopulations contain neurons with both contralateral and ipsilateral choice preferences. On the other hand, the unilateral subpopulations contain neurons with only contralateral choice preference, so we only fit the single accumulator to them. The best initial parameters of the dynamic models were selected through a greedy search approach (See Methods).

**Figure 4.**
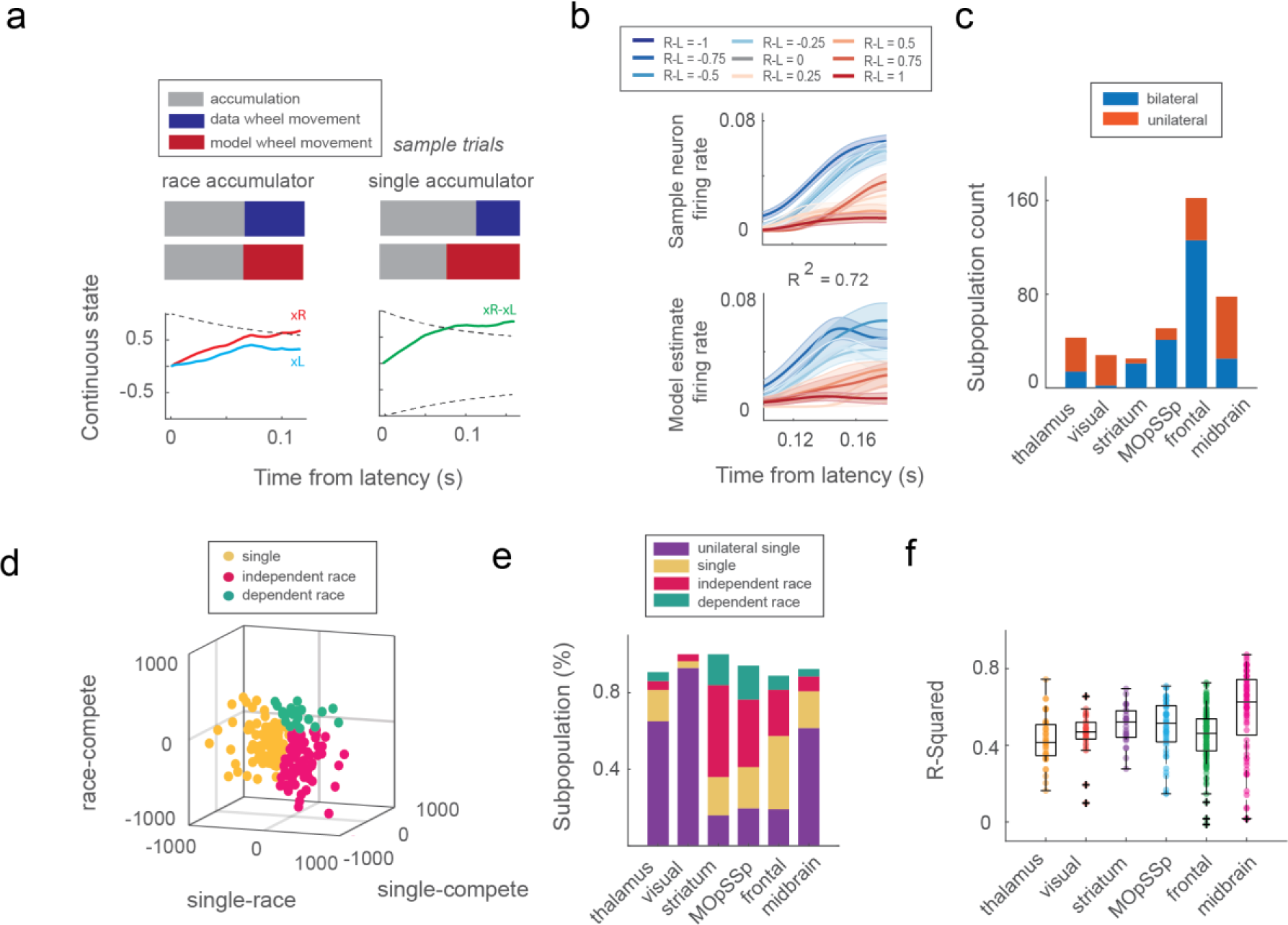
Evaluation of the single and race accumulator models. (a) The discrete state switches to the right choice state when the continuous variable reaches the collapsing boundary. (b) The firing rate of the sample neuron and the fitted single accumulator model. The explained variance (R-squared) between the data and model firing rate is depicted above the figure. The colors indicate the strength of the evidence level. (c) The number of bilateral and unilateral subpopulations within the brain regions. (d) Model comparison using the AIC difference approach. The colors indicate the accumulator type. (e) The percentage of bilateral and unilateral subpopulations preferring single, independent race, and dependent race accumulators. (f) Explained variance (R-squared) values for each brain region’s bilateral and unilateral subpopulations. R-squared values were computed between data and the best model selected using the AIC difference for the bilateral subpopulations. For the unilateral subpopulations, this metric was computed using the single accumulator.

Since we modeled the evidence accumulation phase of the decision-making process, we excluded the neural activity during the visual encoding phase from the accumulator modeling by estimating the accumulation latency using the auROC metric (See Methods). The evolution of the single and independent race variables in sample trials is illustrated in Figure 4a. As shown in this figure, the discrete state switches to the wheel movement state when the continuous variables reach the decision boundary.

The best model for bilateral subpopulations was determined using the AIC difference approach (Figure 4d) (See Methods). We have also computed the explained variance (R-squared) of the models in both bilateral and unilateral subpopulations (Figure 4f, supp-Figure 3). The results demonstrated that bilateral subpopulations within the striatum and MOpSSp regions prefer race accumulators significantly more than the single accumulator (sign test, p-value < 0.05) (supp-Figure 3b). However, we didn’t observe a significant difference between the number of single and rece accumulators for bilateral subpopulations within the frontal area. Moreover, the brain regions which are more unilateral (i.e., visual, thalamus, and midbrain) didn’t produce significant results due to the scarcity of bilateral subpopulations within these regions. Therefore, we also compared the number of single and race accumulators among total subpopulations assuming that unilateral subpopulations could just prefer single accumulators (supp-Figure 3c). As you can see in this figure, the thalamus, visual, and midbrain areas, which are more unilateral, prefer the single accumulator significantly more than race accumulators (sign test, p-value < 0.001). We also observed a significant difference between the number of single and race accumulators within the frontal region (sign test, p-value < 0.001), suggesting that this area prefers the single accumulator more than the race ones.

### Distributed evidence accumulation over multiple timescales

The distributed coding of evidence accumulation across the brain suggests that the accumulation process is happening over multiple timescales, which can be organized hierarchically across the brain (Murray et al., 2014; Pinto et al., 2022). The ability of the brain to function in different timescales stems from the heterogeneity of local microcircuits and their long-range connectivity (Chaudhuri et al., 2015). Here, we examined whether the single and race accumulator models across the brain have distinct properties in terms of the integration timescale. Accordingly, we simulated neurons’ activity within each subpopulation using the preferred accumulator model. The integration timescale was estimated using the combined autocorrelation structure of the simulated neurons’ activity at both the local subpopulation and global population levels within the brain regions (See Methods) (Figure 5a and 5b). The estimated population-level timescale displayed a hierarchical organization across the brain, starting from the visual to the frontal in the cortical regions and the thalamus to the midbrain in the subcortical ones (Figure 5b), which is consistent with previous studies (Chaudhuri et al., 2015; Pinto et al., 2022). The resulting hierarchy demonstrates that thalamic and visual areas integrate the information in a shorter timescale than the midbrain and frontal regions.

**Figure 5.**
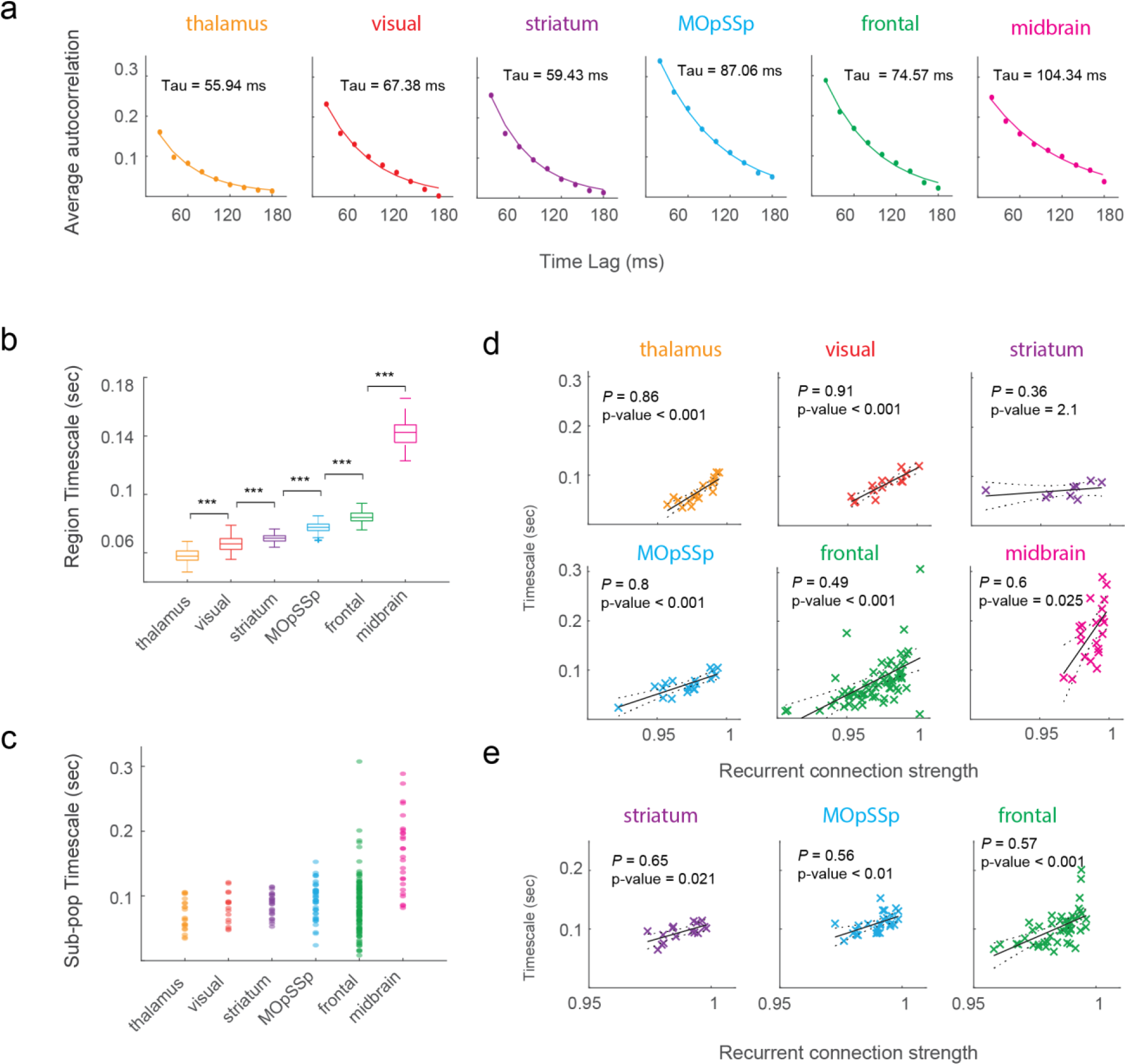
Distribution of the integration timescale across the brain. a) Autocorrelation structure of a simulated subpopulation of neurons is described using the exponential decay function b) Hierarchical organization of the brain areas in terms of the integration timescale. Timescales were estimated using the combined autocorrelations of the sampled subpopulations during a 100-times bootstrapping process. Marker ‘***’ indicates the p-value < 0.001 in the Wilcoxon rank sum test corrected for multiple comparisons. c) Heterogeneity of the subpopulations’ timescale within each brain area. d) Pearson’s correlation between the recurrent connection strength and the integration timescale of single accumulators within each brain area. P-values were corrected by the Bonferroni multiple comparison correction. e) Pearson’s correlation between the average recurrent connection strength of the left and right accumulator variables and the integration timescale of race accumulators within each brain area. P-values were corrected by the Bonferroni multiple comparison correction.

In addition to the hierarchical organization of integration timescale, we also observed a heterogeneity of timescales within each brain area (Figure 5c). We hypothesized the observed diversity of integration timescales could reflect the differences in the accumulator microcircuits. To address this hypothesis, we explored the association between the integration timescale and the recurrent connection strength of the accumulators within each brain area using Pearson’s correlation. The results demonstrated that the recurrent connection strengths of single accumulators were significantly correlated with the integration timescales in most of the regions (Figure 5d). We also examined Pearson’s correlation on the bilateral subpopulations preferring race accumulators by excluding regions with insufficient samples (less than 10 subpopulations) (Figure 5e). The results revealed that the average recurrent connection strengths of the left (*x*_*L*_) and right (*x*_*R*_) accumulators in the race microcircuits (Figure 3b) were significantly correlated with the integration timescales in all the remaining regions. Our findings support the hypothesis that microcircuits with longer integration timescales have larger recurrent connection strength, which is in line with the previous studies (Chaudhuri et al., 2015).

## 3 Discussion

Although previous studies on perceptual decision-making revealed the distribution of decision coding in the mouse brain (Steinmetz et al., 2019), the contribution of these neurons to the evidence accumulation process and the underlying accumulation mechanism remain unclear. Using brain-wide electrophysiological recording in mice (Steinmetz et al., 2019), we showed that evidence accumulation during perceptual decision-making is a distributed process across the brain. We found different cortical and subcortical areas, i.e., visual and frontal cortices, MOp, striatum, midbrain, and thalamus, contain neurons with Drift-Diffusion-Model-like (i.e., evidence-sensitive ramping firing rate) activity. We showed that these regions consist of subpopulations that accumulate evidence through both single and race accumulation mechanisms. We further characterized the accumulation process in terms of the integration timescale. Our findings revealed a hierarchical organization of timescales across the brain, suggesting the existence of evidence accumulation over multiple timescales. In addition, we observed a heterogeneity of timescales within the brain regions, reflecting the diversity of the accumulator’s recurrent connection strength.

The identified brain regions in this study are consistent with and complement the existing findings on the neural substrates of evidence accumulation. Prior work has demonstrated the contribution of a subset of these areas i.e., PPC (Roitman & Shadlen, 2002; Shadlen & Newsome, 2001), FEF (Ding & Gold, 2012; Kim & Shadlen, 1999), striatum (Ding & Gold, 2010), superior colliculus (Horwitz & Newsome, 1999) and FOF (Hanks et al., 2015) in the evidence accumulation process.

The neurons with DDM-like firing rate activity across the brain could integrate the information through single or dual accumulation mechanisms (Bogacz et al., 2006). However, the dual accumulator needs the neural populations supporting each choice alternative. The brain regions we examined contain neurons with both contralateral and ipsilateral choice preferences in the left hemisphere, which were mostly observed in the frontal area. The bilateral behavior of the regions suggested the existence of a dual accumulation mechanism within a single hemisphere, consistent with the previous studies (Mante et al., 2013; Ratcliff et al., 2007; Wong et al., 2007). We tried to investigate whether DDM-like neurons in the brain were best represented using single or dual accumulators.

Our results revealed that bilateral subpopulations within the striatum and MOpSSp strongly prefer race accumulators more than single ones. However, exploring the accumulator preferences among the combined unilateral and bilateral subpopulations demonstrated that the visual, thalamus, and midbrain regions strongly prefer the single accumulator. This may be due to the unilateral nature of these brain regions. However, despite the bilateral nature of the frontal area, the number of subpopulations with single accumulation preferences is higher than the ones preferring dual accumulators. This may be due to the single-hemisphere neural recording.

We sought to address whether the distributed nature of evidence accumulation processes was related to how neurons in different brain regions represent information at different timescales. The estimated accumulator’s integration timescale at the population level revealed hierarchical organization across the brain regions. According to this hierarchy, the integration timescale increases from visual to frontal in the cortical regions and from the thalamus to the midbrain in the subcortical ones, consistent with the previous studies (Chaudhuri et al., 2015; Honey et al., 2012; Pinto et al., 2022). Our findings lend further support to previous claims that evidence accumulation is happening over multiple timescales, and different brain areas in humans, primates, and rodents display a hierarchical organization in terms of their timescale (Chaudhuri et al., 2015; Demirtaş et al., 2019; Gao et al., 2020; Honey et al., 2012; Imani et al., 2023; Murray et al., 2014; Pinto et al., 2022; Rossi-Pool et al., 2021). We extend this literature (e.g., for most recent findings using calcium imaging data in cortical regions see Pinto et al. 2022) by providing evidence from the analysis of electrophysiological data across the whole mouse brain. This hierarchical organization could be an essential component of the distributed evidence accumulation process across the brain (Pinto et al., 2022), which may be due to the variability in the level of recurrent excitation connections within areas (Chen et al., 2015; Gao et al., 2020), and their long-range connectivity profile (Chaudhuri et al., 2015). The hierarchical organization of the brain areas in terms of the integration timescale also suggests that the inactivation of brain areas across the cortical hierarchy could affect the performance of the decision-making process at different timescales (Pinto et al., 2022; Zatka-Haas et al., 2021). In addition to the variability of timescale across the brain, we observed heterogeneity of timescale within each brain area. Our findings suggest that this heterogeneity may arise from the variation in the local accumulation microcircuits. Such that, accumulators with longer integration timescales have higher recurrent connection strength, which is consistent with the previous studies (Chaudhuri et al., 2015).

In summary, we have investigated the neural correlate of evidence accumulation across the brain. We identified that DDM-like neurons are distributed across the brain, which can integrate information through single or dual accumulation mechanisms. These accumulator circuits were characterized using distinct integration timescales which were organized hierarchically across the brain. Our findings support the hypothesis that evidence accumulation is a distributed process over multiple timescales. Moreover, we observed a heterogeneity of integration timescales within each brain area suggesting a diversity of accumulator microcircuit parameters.

## 4 Materials and Methods

### 4-1 Behavioral task

We used a publicly available dataset published recently by (Steinmetz et al., 2019). The dataset comprises behavioral and physiological data from ten mice over 39 sessions on a two-alternative unforced choice task. Mice sit on a plastic apparatus with their forepaws on a rotating wheel, surrounded by three computer monitors. At each trial that was started by briefly holding the wheel, visual stimuli (Gabor patch with sigma 9 and 45° direction) with four grading levels were displayed on the right, left, both, or neither screen (Figure 1a). The stimulus was presented in the mouse’s central monocular zones, and the animal did not need to move its head to perceive it.

Mice earned water by turning the wheel to move the stimulus with the highest contrast to the center of the screen or by not turning the wheel if neither stimulus was displayed. Otherwise, they received a white noise sound for one second to indicate an improper wheel movement. Therefore, three types of trial outcomes (right turn, left turn, and no turn) leads to reward. After the stimulus presentation, a random delay interval of 0.5–1.2s was considered, during which the mouse could freely turn the wheel without incentive. At the end of the interval, an auditory tone cue (8 kHz pure tone for 0.2s) was played, at which point the visual stimulus position became coupled with the wheel movement.

### 4-2 Neural recording

Recordings were made in the left hemisphere using the Neuropixel electrode arrays from approximately 30,000 neurons in 42 brain areas in 39 sessions. Using the Neuropixel probes with the ability to record from multiple brain regions produced data simultaneously recorded from several regions in each session. The neural activity of the regions was divided into seven main groups according to the Allen Common Coordinate Framework (CCF) atlas (Wang et al., 2020) (Figure 1b). We performed all the analyses on these groups of regions.

### 4-3 Single neuron decoding analysis

We performed the single neuron decoding using the area under the receiver operating characteristic (auROC) analysis. The auROC metric was initially employed to measure the neuron’s choice probability based on the Mann–Whitney U statistic (Britten et al., 1996). Using this nonparametric statistical method, we can measure the differences between spike count distributions of two conditions (or behavioral outputs) to examine whether the neuron’s firing rate is significantly greater than the other condition. According tothe task design, the stimulus and choice encoding are highly correlated and cause the decoding analysis. To overcome this limitation, we used combined condition auROC analysis to compute stimulus selectivity, choice probability, detect probability, and evidence selectivity. The trials were then divided into different groups according to the task conditions, and the weighted average of the auROC values across conditions was considered the final decoding result. For this analysis, the neuron’s spikes were binned at 0.005s and smoothed using a causal half-Gaussian kernel with a standard deviation of 0.02s. We also z-scored the firing rate of the neurons by subtracting the mean and dividing by the standard deviation calculated during the baseline period (-0.9s to -0.1s, stimulus aligned) across all trials.

### 4-3-1 Stimulus selectivity

We computed the contra stimulus selectivity using the combined condition auROC metric. Accordingly, the trials were divided into 12 groups based on the different choice alternatives (right, left, NoGo) and stimulus contrast levels (0, 0.25, 0.5, 1) presented on the left screen. We then applied the Mann–Whitney U statistic to measure the stimulus selectivity by comparing the spike counts of a neuron within the trials with the right stimulus higher than zero with the trials having the right stimulus equal to zero. The final stimulus selectivity was measured using the weighted average across 12 conditions.

### 4-3-2 Choice Probability

Using the combined condition auROC statistic, we tested whether the neurons encode the choice. To compensate for the effect of the stimulus conditions, we divided the trials into 12 groups based on different combinations of right and left stimulus contrast levels, ignoring equal contrast conditions. Within each condition, we used the Mann–Whitney U statistic to compare the spike count of the trials with right/left choice with another choice in a window ranging from -0.3s to 0.1s (aligned with wheel movement). A weighted average was then utilized to compute the final choice probability over different conditions. The absolute deviation of auROC from the chance level was considered as the choice selectivity: *CP* = |*auROC* − 0.5|.

### 4-3-3 Detect probability

We also measured how well the neural activity encodes whether or not the animal turned the wheel correctly and referred to this measurement as ‘Detect probability’ (Hashemi et al., 2018). Accordingly, the trials were divided into 12 groups based on the different combinations of the right and left stimulus contrast levels, excluding the conditions with equal contrast levels. We then measured whether the Hit (correctly turning the wheel) trials had greater neural activity than the Missed trials using the Mann–Whitney U statistic during the stimulus epoch (-0.1s to 0.3s). The level of selectivity for this measurement was calculated as the deviation of auROC from the chance level: *DP* = *auROC* − 0.5.

### 4-3-4 Evidence selectivity

We measured how a neuron can encode the evidence (difference of right and left stimulus contrast levels) and defined it as ‘Evidence selectivity’. The trials were divided into nine groups according to the number of evidence levels (ranging from -1 to 1 with a step size of 0.25). We then tested to see whether the group of trials with the higher evidence level had greater neural activity than all those groups with lower evidence. Final evidence selectivity was calculated by taking the weighted average of auROC values across eight group comparisons. Absolute deviation of auROC from the chance level was considered as the measure of evidence selectivity: *ES* = |*auROC* − 0.5|.

### 4-3-5 Significant auROC selectivity

We also performed the auROC analysis on the shuffled trial labels to identify significantly selective neurons. We created the distribution of the auROC on the shuffled trials by repeating the shuffling process 100 times. The selectivity of a neuron at time *t* was considered significant if the value of the true auROC was outside the confidence interval of the shuffled auROC values. We restricted our analysis to the time points with at least two significant neighbors to correct the multiple comparisons.

### 4-4 Neuron latency

Evidence accumulation usually starts after a latency, mainly related to the visual encoding state (Roitman & Shadlen, 2002). In this study, we restricted the evidence accumulation analysis to the neural activity within the window starting at the end of latency until 50ms after wheel movement. We used auROC analysis to compute latency which appears like the time of significant change in neural activity compared with the baseline activity.

Accordingly, the spike counts of the Go trials having reaction times within the range (0.15s to 0.5s) were smoothed using a causal boxcar filter of size 100ms during (-0.5s to 0.5s) aligned to stimulus onset. We then computed the average firing rate across neurons within each trial, followed by the Mann–Whitney U statistic to compare the neural activity of trials within the stimulus (0s to 0.5s) and a point in the baseline (-0.1s) epochs. The significance level (*p*-value < 0.05) was employed to detect the samples with significant neural activity changes. We restricted our analysis to the significant points with at least two significant neighbors to correct for multiple comparisons. The latency was then selected as the first time point with significant changes in neural activity.

### 4-5 Demixed principal component analysis (dPCA)

Most neurons, especially in the higher cortical areas, encode different types of task information and display a mixed selectivity (Kobak et al., 2016). This complexity in response selectivity of the neurons can conceal their expressed information. To overcome this limitation, we exploited the advantage of demixed principal component analysis (dPCA) to decompose the population neural activity into a few latent components, each capturing a specific aspect of the task (Kobak et al., 2016). The resulting dPCA subspace captures most variation in the data and decouples different task-related components.

According to the dPCA analysis, we prepared a matrix *X*_*N*×4×2×*T*_ containing marginalized activity of N neurons over four stimulus contrast levels (1, 0.5, 0.25, 0) and two decision alternatives (Hit and Missed) during T time points within the stimulus epoch (-0.1s to 0.3s). We excluded the neurons having an average spike count lower than 1Hz during the reaction time boundary (from stimulus onset until wheel movement). We first divided trials into four groups to construct the matrix based on the contralateral stimulus contrast levels. Within each group, the average trial activity of each neuron was then computed based on whether or not the animal turned the wheel. The dPCA algorithm was applied to the neural population matrix to construct a latent subspace with 20 task-related principal components. The resulting components characterized the decision (*X*_*dt*_), stimulus (*X*_*st*_), stimulus-decision interaction (*X*_*sdt*_), and condition-independent (*X*_*t*_) information by estimating task-specific decoders *F*_*α*_ and encoders *D*_*α*_ using the following loss function (Kobak et al., 2016):

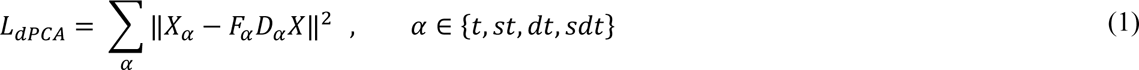

We computed the explained variances *R*^2^ (R-squared) of the neurons by projecting the neural activity to the task-specific principal components using the decoder matrices *D*_*α*_ and reconstructing the neural activity with the decoder matrices *F*_*α*_ as follows:

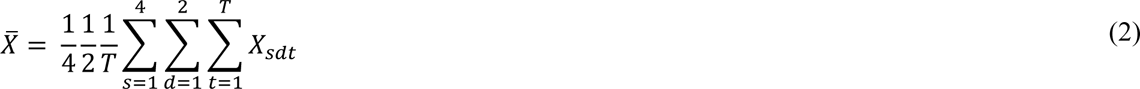

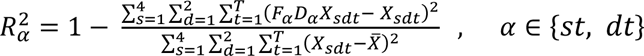

We then separated the neurons into stimulus, decision, and interaction groups within each brain region using their task-related R-squared values (*R*^2^, *R*^2^) and fuzzy C-means clustering algorithm (Bezdek, 2013). We excluded the neurons within the stimulus and interaction clusters from further analysis.

### 4-6 Integration timescale

We measured the integration timescale of the subpopulations using the spike count autocorrelation structure of the simulated neural activity. Accordingly, we simulated fixed-length trials of duration 200ms for each subpopulation using the preferred (single or race) accumulator model during a 50-times sampling process. We then estimated the timescale of simulated neurons within each sample set of trials and considered the average timescale across the 50 samples as the final timescale for the subpopulation. To estimate the timescale of simulated neurons at each sampling iteration, we computed the Pearson’s correlation of binned spike counts between each pair of time bins *iΔ* and *jΔ* (*j* > *i*, *Δ* = 0.025*s*) across trials. The resulting autocorrelation values follow an exponential decay which can be explained using the following equation (Murray et al., 2014):

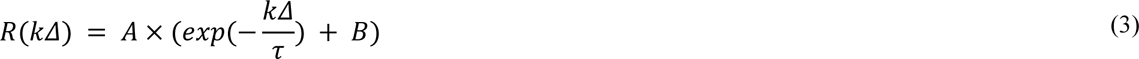

[where A is the amplitude, B indicates the contribution of timescales longer than the observation window, *kΔ* is the time lag, and *τ* denotes the timescale.

We fitted equation (3) to the combined autocorrelation structure of the simulated neurons within each subpopulation using the Levenberg-Marquardt method. The time lag with the greatest autocorrelation reduction was selected as the starting point for overcoming the negative adaptation (Murray et al., 2014). We tried five different initial parameter values to select the best model having the lowest mean square error (MSE) value. We eventually computed the average of timescales across 50 sets of simulated trials.

Similarly, the global population-level timescale of each brain region was estimated based on the combined autocorrelation structure of the simulated neurons. We bootstrapped subpopulations within each brain area 100 times to compute the confidence interval of the population-level timescales. We further applied a Wilcoxon rank sum test on the bootstrapped samples to test for significant differences between regions.

### 4-7 Recurrent switching linear dynamical system (rSLDS)

We employed a general framework proposed by (Zoltowski et al., 2020) for modeling the evidence accumulation process. Different evidence accumulation models are formulated in this framework as a recurrent switching linear dynamical system (rSLDS). The rSLDS contains multiple discrete states *z*_*t*_ ∈ {1, …, *K*} and each state is associated with specific linear dynamics (Figure 3a) as follows:

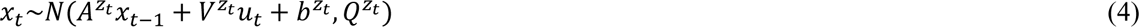

where *x*_*t*_ ∈ *R*^*D*^ is the continuous state, *u*_*t*_ ∈ *R*^*M*^ represents input streams, *Q*^*z*_*t*_^ ∈ *R*^*D*×*D*^ is the noise covariance matrix, and the matrices *A*^*z*_*t*_^ ∈ *R*^*D*×*D*^ and *V*^*z*^_*t*_ ∈ *R*^*D*×*M*^, and vector *b*^*z*^_*t*_ ∈ *R*^*D*^ denote the state-specific dynamic parameters. Transition probabilities between discrete states are parameterized as follows:

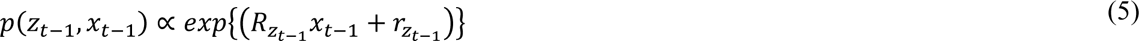

where *R*_*z*_*t*−1__ ∈ *R*^*K*×*D*^ and *r*_*z*_*t*-1__ ∈ *R*^*K*^ parameterize the influence of the continuous state on the discrete state transitions. The observation model was used to map the continuous latent variables *x*_*t*_ into the overserved variable *y*_*t*_ using the Poisson distribution of a generalized linear model as follows:

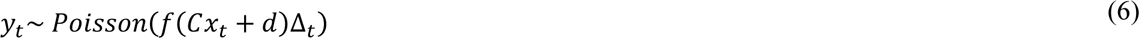

where *f*(*x*) = *log*(1 + *exp* (*x*)) is the Softplus function, and Δ_*t*_ denotes the size of time bins. The weight parameter *C ∈ R*^*D*×*N*^ was used to map the latent variable *x*_*t*_ ∈ *R*^*D*^ into the activity of N neurons *y*_*t*_. However, the offset parameter *d ∈ R*^*D*^ in the observation model is shared across the neurons.

#### 4-7-1 Single accumulator

A single accumulator model, which is commonly referred to as the drift-diffusion model (DDM), is described with a single decision variable that accumulates the differences in the input streams (Bogacz et al., 2006). This accumulation mechanism has two decision boundaries, one for each choice alternative. When the decision variable reaches one of the boundaries, the decision is made.

To reformulate the rSLDS framework to a single accumulator, we considered three discrete states for the accumulation (*z*_*t*_ = *acc*) phase, right wheel movement (*z*_*t*_ = *rwm*), and left wheel movement (*z*_*t*_ = *lwm*) (Figure 3c). During the evidence accumulation state, the one-dimensional continuous variable *x*_*t*_ ∈ *R*^1^ accumulates the differences between right and left input streams *u*_*t*_ ∈ *R*^1^. The state transition was also parameterized such that when the continuous variable *x*_*t*_ reaches one of the decision boundaries (±B), the discrete state switches from the accumulation state (*z*_*t*_ = *acc*) to the right wheel movement (*z*_*t*_ = *rwm*) or left wheel movement (*z*_*t*_ = *lwm*) states. Therefore, the transition parameters were set as follows:

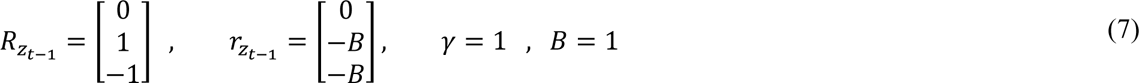

According to the settings, increasing the value of *x*_*t*_ toward B leads to an increase in the probability of transition from the *z*_*t*_ = *acc* to *z*_*t*_ = *rwm*. On the other hand, decreasing the value of *x*_*t*_ toward -B, increases the probability of transition to *z*_*t*_ = *lwm*.

In equation (4), the term *A* ∈ *R*, denotes the recurrent connection strength, and the term *V* ∈ *R* determines the weight of the received input stream (Figure 3b). We excluded the term *b* ∈ *R* from our analysis. In the accumulation state (*z*_*t*_ = *acc*), we only trained the term *A*^*acc*^ and the noise *Q*^*acc*^, and set the term *V*^*acc*^ constant. In other states (*z*_*t*_ = *rwm*, *lwm*), we just trained the noise variance Q and considered *A*^*rwm*^ = *A*^*lwm*^ = 1 and *V*^*rwm*^ = *V*^*lwm*^ = 0. We tried different initial values for the parameters *A*^*acc*^ ∈ {0.95, 1}, *V*^*acc*^ ∈ {0.01, 0.02, 0.03, 0.04, 0.05}, *Q* ∈ {0.005, 0.01} to select the best combination of parameters that produced maximum log-likelihood.

#### 4-7-2 Independent race accumulator

An independent race accumulator model contains two integrators that accumulate the relative or absolute input streams supporting each choice alternative (Bogacz et al., 2006). In this accumulation mechanism, a decision is made favoring the integrator that reaches the decision boundary sooner. To reformulate the rSLDS into an independent race accumulator mechanism, we considered a two-dimensional continuous variable *x*_*t*_ ∈ *R*^2^ for two accumulators. These variables accumulated the absolute right/left input streams *u*_*t*_ ∈ *R*^2^ independently. We set the parameters of the dynamic model (*A*_*acc*_, *V*_*acc*_, *Q*_*acc*_) to be diagonal such that the decision variables integrate the input streams independently. Similar to the single accumulator, we considered three discrete states for the accumulation (*z*_*t*_ = *acc*) phase, right wheel movement (*z*_*t*_ = *rwm*), and left wheel movement (*z*_*t*_ = *lwm*) (Figure 3c).

We also set the transition parameters such that the probability of switching from the accumulation state to one of the wheel movement states increases by approaching *x*_*t*_ to decision boundary *B*. Accordingly, the transition parameters were set as follows:

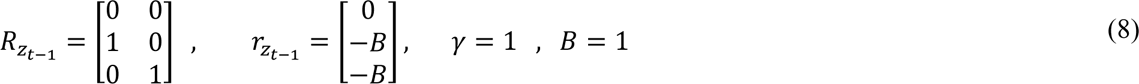

In equation (4), on-diagonal values in matrix *A* ∈ *R*^2^ determines the excitatory connection strength for two accumulators. The on-diagonal values in matrix *V* ∈ *R*^2^ also denotes the weight of the received input stream for each accumulator variable (Figure 3b). Similar to the single accumulator mechanism, we excluded term *b* from our analysis. In the accumulation state (*z*_*t*_ = *acc*), we only trained matrices *A*^*acc*^ and noise *Q*^*acc*^, and set the matrix *V*^*acc*^ constant. In other states (*z*_*t*_ = *rwm*, *lwm*), we just trained the noise covariance matrix Q and considered *A*^*rwm*^ = *A*^*lwm*^ = 1 and *V*^*rwm*^ = *V*^*lwm*^ = 0_2,2_. We tried different initial values for the on-diagonal values of matrices *A*^*acc*^ ∈ {0.95, 1}, *V*^*acc*^ ∈ {0.01, 0.02, 0.03}, *Q* ∈ {0.005, 0.01} to select the best combinations of parameters that produced maximum log-likelihood.

#### 4-7-3 Dependent race accumulator

Dependent race models are a more general form of dual accumulators containing mutual (Machens et al., 2005; Usher & McClelland, 2001; Wong & Wang, 2006) and feedforward connections (Palmeri et al., 2015; Purcell et al., 2010). In these models, each decision variable accumulates input streams supporting each choice alternative and the decision is made favoring the integrator that reaches the respective decision boundary sooner.

Similar to the independent race model, we consider a two-dimensional continuous variable *x*_*t*_ ∈ *R*^2^ to reformulate rSLDS into the dependent race accumulator. To model the mutual and feedforward connections, we considered parameters in the dynamic model (*A*_*acc*_, *V*_*acc*_, *Q*_*acc*_) to be fully connected rather than diagonal. Due to the negative and positive decision boundaries in this model, we considered five discrete states for the accumulation (*z*_*t*_ = *acc*) phase, positive/negative right wheel movement (*z*_*t*_ = p*rwm*, *z*_*t*_ = n*rwm*), and positive/negative left wheel movement (*z*_*t*_ = p*lwm*, *z*_*t*_ = n*lwm*).

The transition parameters are set such that the probability of switching from the accumulation state to the right or left wheel movement states increases by approaching *x*_*t*_ to each of the decision boundaries ±*B*. Accordingly, the transition parameters were set as follows:

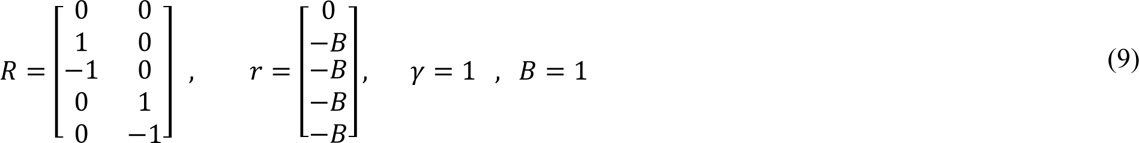

In the accumulation state (*z*_*t*_ = *acc*), we only trained matrices *A*^*acc*^ and noise *Q*^*acc*^, and set the matrix *V*^*acc*^ constant. In the other four states, we just trained the noise covariance matrices Q and considered matrices A and V to be the identity and null matrices, respectively. We tried different initial values for the on-diagonal parameters *A*^*acc*^ ∈ {0.95, 1}, *V*^*acc*^ ∈ {0.01, 0.02, 0.03}, *Q* ∈ {0.01} and off-diagonal parameters *A*^*acc*^ ∈ {−0.05}, *V*^*acc*^ ∈ {−0.01, − 0.02, − 0.03} to select the best combinations of parameters that produced maximum log-likelihood.

#### 4-7-4 Collapsing boundary

In the accumulators with the collapsing boundary, less evidence is required to reach the boundary as time passes so that the boundaries collapse toward the center (Figure 3d). This mechanism is much like the urgency signal, magnifying the evidence as time passes (Ratcliff et al., 2016). Besides the constant decision boundaries, we also evaluated the collapsing boundary in single and dual accumulators.

In the rSLDS framework, we can reformulate equation (5) to implement the linear collapsing boundary for a single accumulator as follows (Zoltowski et al., 2020):

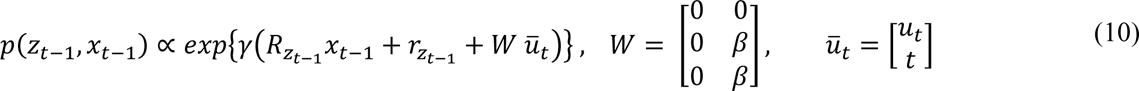

Where *W* <u>*u*</u>_*t*_ is the linear function of time points *t* and vector <u>*u*</u>_*t*_ contains the input streams and the current time. This equation describes a linear collapsing boundary with the rate of *β*. We need to add another column to the matrix V in equation (4) and set it to zero with this new formulation. We further modified equation (9) to formulate a nonlinear collapsing boundary for a single accumulator as follows:

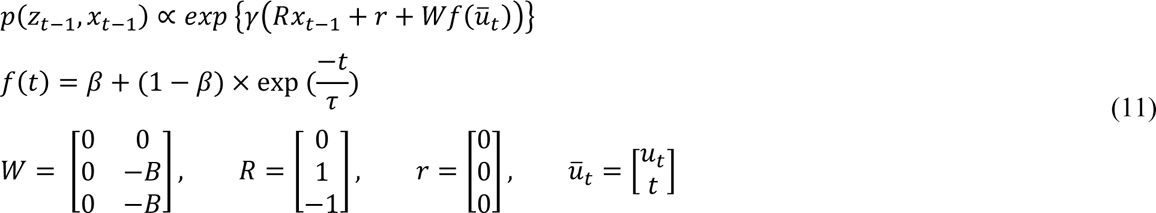

Where *β* denotes the boundary offset and *τ* describes the decay rate of the exponential function. We can control the collapsing rate with these two parameters (Figure 3d). To implement the collapsing boundary for the independent and dependent race accumulators, we set the parameters of the transition model as equations (12) and (13) respectively:

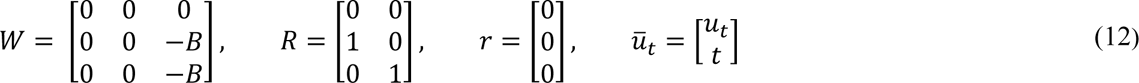

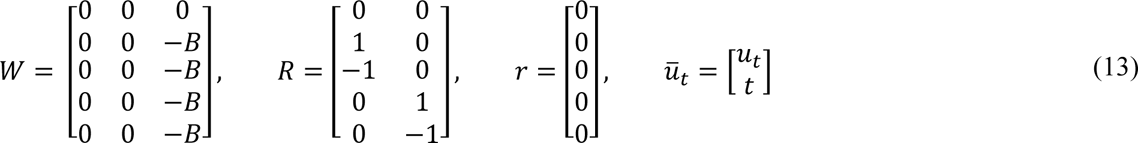

We tried different initial values for *β* ∈ {0.3, 0.4, 0.5} and *τ* ∈ {50, 100} to select the best parameter which leads to the maximum log-likelihood.

#### 4-7-5 Model fitting

We fit the accumulator models to the subpopulations of neurons within the brain regions at each session. Accordingly, subpopulations were generated by sampling four DDM-like neurons without replacement within each brain area. To improve the performance of modeling, we excluded trials according to the stimulus and reaction time criteria. Accordingly, trials with equal contrast levels (Right = Left) were excluded due to the random behavioral output of mice during these trials. We further focused our analysis on the trials with reaction times longer than 150ms and shorter than 500ms.

To model the evidence accumulation process, we did not consider fixed-length trials. Given that the perceptual decision-making process comprises different cognitive stages (visual encoding, evidence accumulation, and action execution) (Mazurek et al., 2003), we excluded the neural activity corresponding to the visual encoding phase (Roitman & Shadlen, 2002). The remaining samples before wheel movement are considered as the evidence accumulation phase. We also included the neural activity from the 50ms post-wheel movement period. This is because of considering multiple discrete states (i.e., accumulation and Right/Left wheel movement phases) to reformulate the recurrent switching linear dynamical system (rSLDS) into different accumulators. According to these settings, the continuous variables evolve in the accumulation state and switch to the Right/Left wheel movement state by reaching the corresponding decision boundary.

Zoltowski et al., 2020 introduced a variational Laplace-EM algorithm to estimate the model parameters. Briefly, the posterior over the discrete and continuous states were calculated using variational and Laplace approximations. The model parameters were also updated by sampling from the discrete and continuous posteriors followed by an Expectation-Maximization (EM) approach (Zoltowski et al., 2020).

#### 4-7-6 Model goodness of fit

##### 4-7-6-1 Akaike Information Criterion (AIC)

We compared the model fitting to the data using the Akaike Information Criterion (AIC) goodness of fit, which is defined as follows (Anderson & Burnham, 2004):

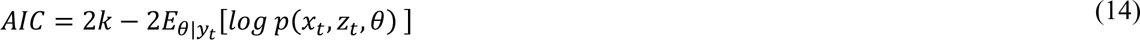

Where *k* is the number of free parameters in the model and the expectation term *E*_θ|*y*_*t* can be estimated by sampling the fitted model 100 times as follows:

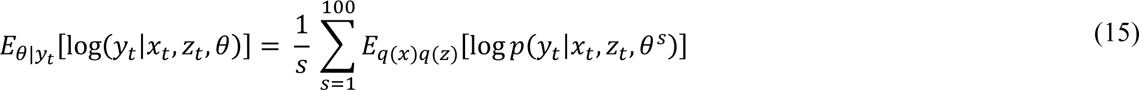

where θ^*s*^ denotes sampling of the model using trained parameters. To compute the log-likelihood, we need to marginalize the hidden variables *x* and *z*. Accordingly, we sampled from the estimated posterior probabilities (*q*(*x*) and *q*(*z*)) to compute the sample-based expectation over these two variables (Zoltowski et al., 2020). AIC measurement contains a penalty term for the number of parameters which is a correction for how much the model with *k* parameters will increase the log-likelihood.

##### 4-7-6-2 R-Squared

We also measured how well a model can explain the data using the R-Squared explained variance. Accordingly, we simulated the spike counts from each model 100 times for each trial. The firing rate of the real and simulated spike counts of subpopulations was computed using a causal boxcar filter of size 50ms, and the average firing rate of trials within each evidence level (Right contrast level-Left contrast level) was computed. We then used the R-Squared explained variance metric on the subpopulations as follows (Latimer et al., 2015):

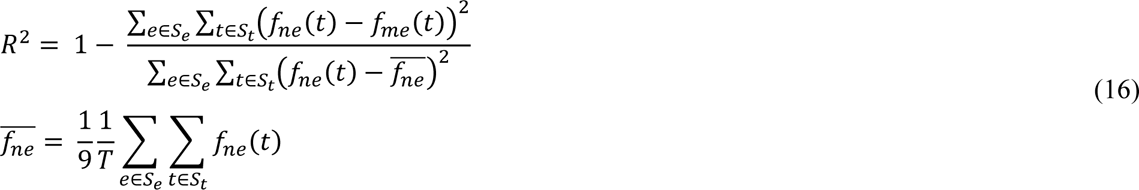

where *e* is the evidence level from the set of evidence *S*_*e*_, terms *f*_*ne*_ and *f*_*me*_ represent the average firing rate of the data and simulated spike counts across the trials with evidence level *e,* respectively. Set *S*_*t*_ denotes the time points within the window from the latency until the median reaction time of the session. The Term <u>*f*_*ne*_</u> is the average firing rate of the data over all time points and coherence levels. *R*^2^ = 1 demonstrates that the model firing rate perfectly matches the data, and lower values correspond to the worst fit.

#### 4-7-7 Model comparison

The preferred accumulator type among the single and race accumulators is selected using the AIC difference approach. According to this approach, the AIC values are rescaled as follows:

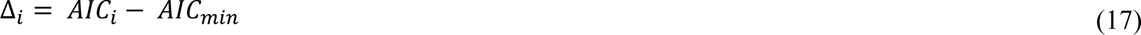

Where *A*1*C*_*min*_ is the minimum of AIC values among the single and race accumulators for a specific subpopulation. According to this transformation, the best model have Δ_*i*_ = 0 and other models have positive Δ_*i*_ values. To select the best model we set the supporting threshold of 10 (Anderson & Burnham, 2004; Latimer et al., 2015). Accordingly, we excluded subpopulations having more than one model with an AIC difference Δ_*i*_ < 10 from our further analysis.

### 4-8 Data processing

All analyses were carried out using customized MATLAB and Python code. Statistical analyses and fuzzy C-means clustering were performed using MATLAB toolboxes. Decomposing neural activity into different task-related variables was carried out using the open-source dPCA toolbox (Kobak et al., 2016). The accumulator analysis was performed using the recurrent switching linear dynamical system (rSLDS) toolbox (Zoltowski et al., 2020), which was customized by the authors.

## Supplemental information

**Supp-Figure 1.**
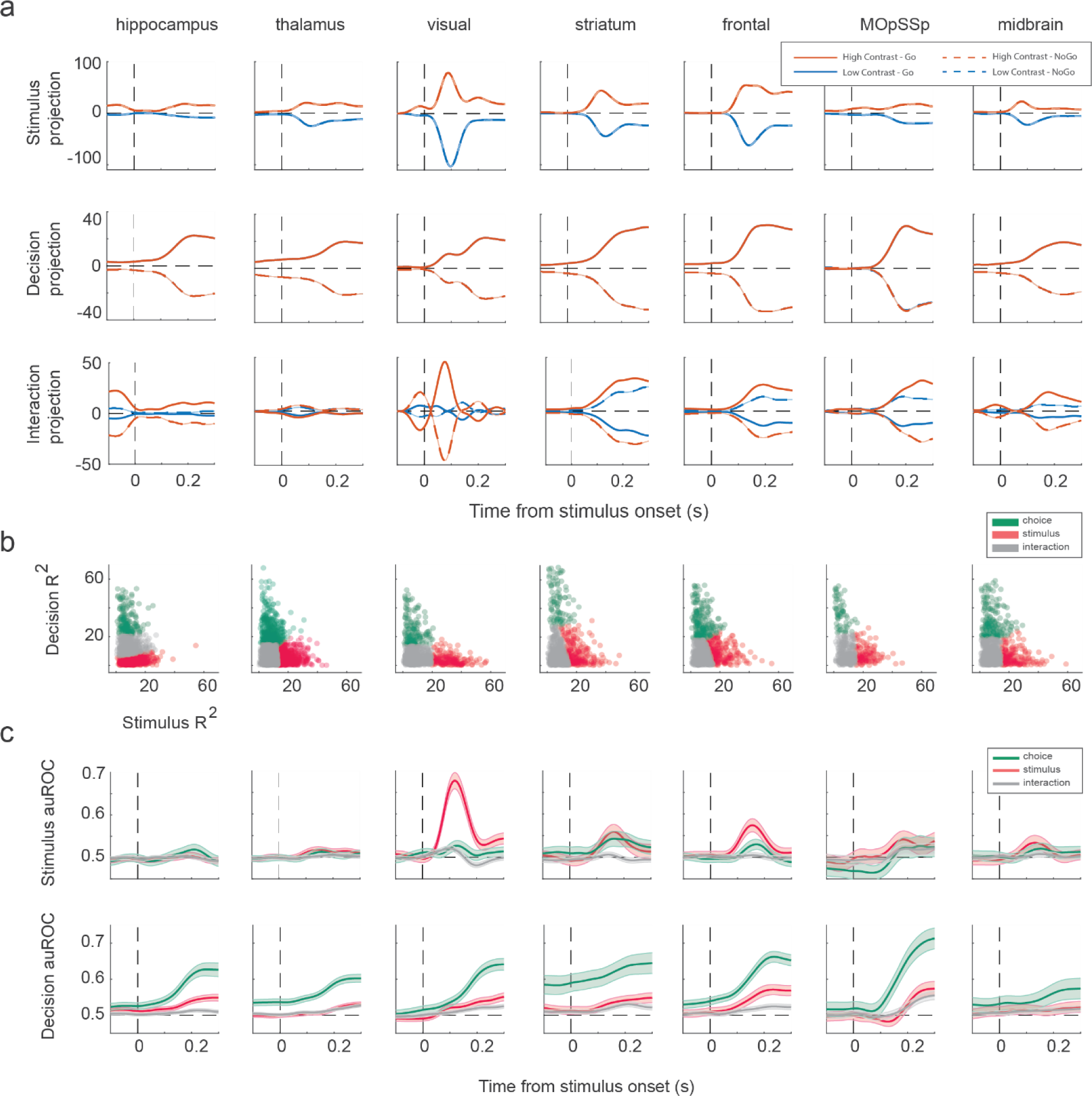
Separating neurons into the decision-selective and the stimulus-selective neurons. (a) Projection of the population neural activity into task-related components. (b) Clustering the neurons based on their stimulus-related and decision-related R-squared values. (c) Performance of the stimulus and decision decoding using each group of neurons (stimulus, decision, and interaction). Shaded areas represent the 95% confidence interval.

**Supp-Figure 2.**
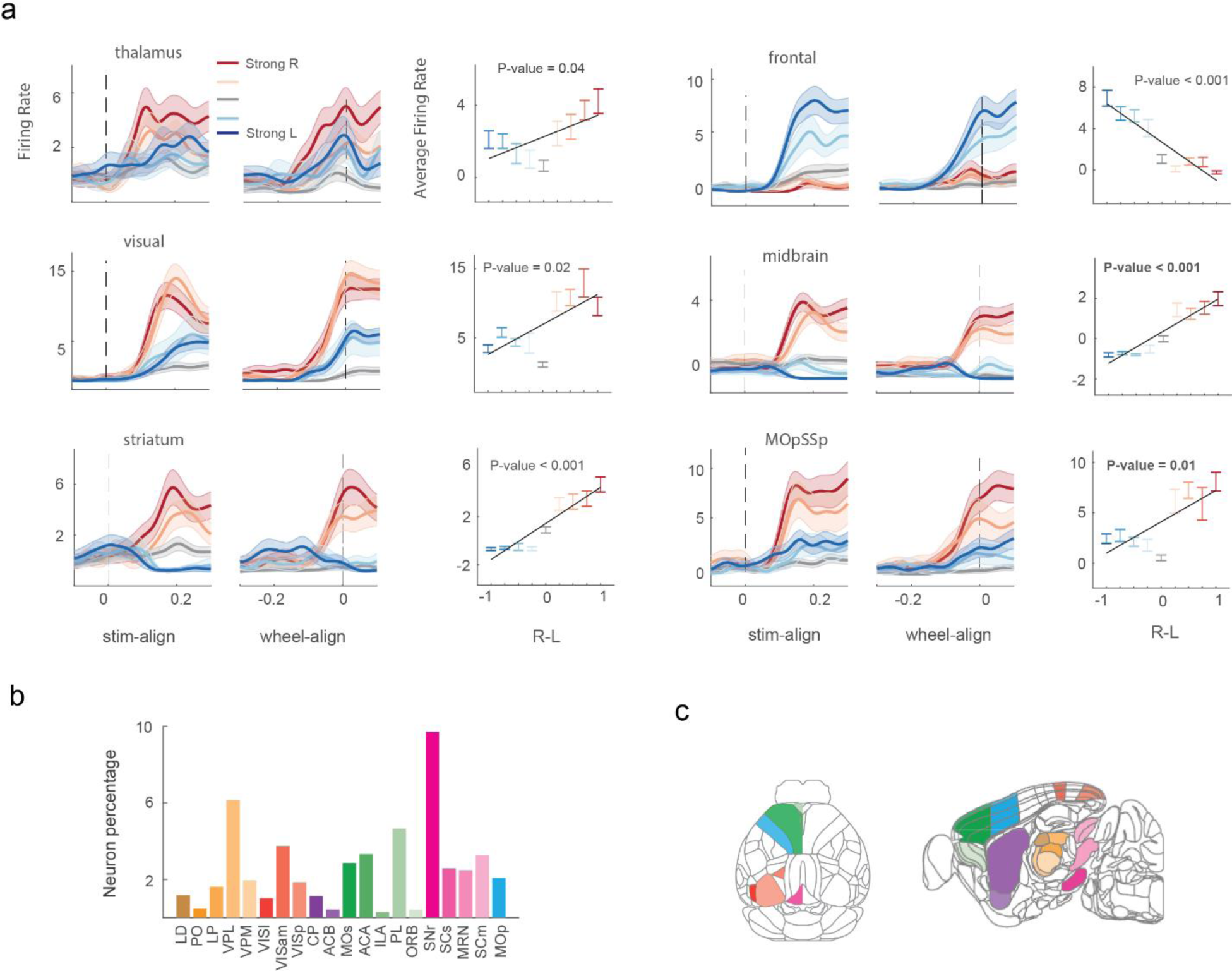
Distribution of DDM-like neurons across the brain. (a) Sample DDM-like neurons. The left panel represents the average firing rate activity of the neuron across trials with a specific evidence level. The strength of the color indicates the strength of the evidence level. Shaded areas represent the confidence interval. The right panels indicate the linear relationship between the average firing rate and the evidence levels using the general linear model. The error bars indicate the 95% confidence interval. (b) The number of DDM-like neurons across different brain areas. (c) Distribution of DDM-like across the brain.

**Supp-Figure 3.**
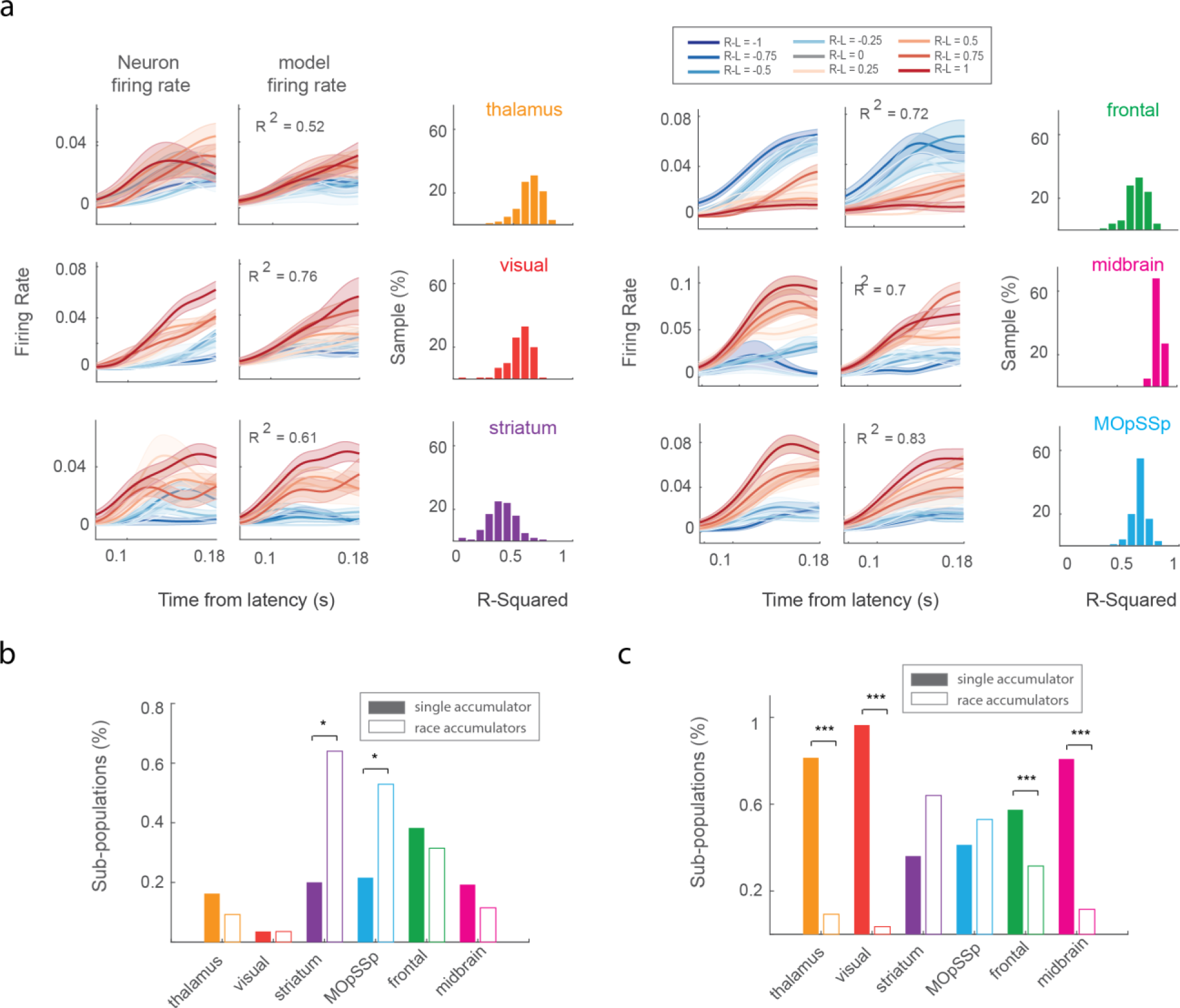
Results of the accumulator fitting. Data and model firing rate of sample neurons and their corresponding explained variance (R-Squared) value. The distribution of R-squared values for each neuron was generated by sampling the accumulator model 100 times. The curves represent the average firing rate activity of the neuron across trials with a specific evidence level. The strength of the color indicates the strength of the evidence level. Shaded areas represent the confidence interval. (b) The proportion of bilateral subpopulations preferring single and race accumulators. Marker ‘*’ indicates the p-value < 0.05 in the sign test. (c) The percentage of the single and race accumulators among the combination of unilateral and bilateral subpopulations. Marker ‘***’ represents the p-value < 0.001 in the sign test. P-values were corrected by the Bonferroni multiple comparison correction.

## Data availability

All neural and behavioral data analyzed in this study are available at https://figshare.com/articles/steinmetz/9598406.

## References

Anderson, D., & Burnham, K. (2004). Model selection and multi-model inference. Second. NY: Springer-Verlag, 63(2020), 10. https://doi.org/10.1007/b97636

Bezdek, J. C. (2013). Pattern recognition with fuzzy objective function algorithms. Springer Science & Business Media. https://doi.org/10.1007/978-1-4757-0450-1

Bogacz, R., Brown, E., Moehlis, J., Holmes, P., & Cohen, J. D. (2006). The physics of optimal decision making: a formal analysis of models of performance in two-alternative forced-choice tasks. Psychological review, 113(4), 700. https://doi.org/10.1037/0033-295X.113.4.700.

Britten, K. H., Newsome, W. T., Shadlen, M. N., Celebrini, S., & Movshon, J. A. (1996). A relationship between behavioral choice and the visual responses of neurons in macaque MT. Visual neuroscience, 13(1), 87–100. https://doi.org/10.1017/s095252380000715x

Chaudhuri, R., Knoblauch, K., Gariel, M.-A., Kennedy, H., & Wang, X.-J. (2015). A Large-Scale Circuit Mechanism for Hierarchical Dynamical Processing in the Primate Cortex. Neuron, 88(2), 419–431. https://doi.org/10.1016/j.neuron.2015.09.008

Chen, J., Hasson, U., & Honey, C. J. (2015). Processing timescales as an organizing principle for primate cortex. Neuron, 88(2), 244–246. https://doi.org/10.1016/j.neuron.2015.10.010

Demirtaş, M., Burt, J. B., Helmer, M., Ji, J. L., Adkinson, B. D., Glasser, M. F., Van Essen, D. C., Sotiropoulos, S. N., Anticevic, A., & Murray, J. D. (2019). Hierarchical heterogeneity across human cortex shapes large-scale neural dynamics. Neuron, 101(6), 1181–1194. e1113. https://doi.org/10.1016/j.neuron.2019.01.017

Ding, L., & Gold, J. I. (2010). Caudate encodes multiple computations for perceptual decisions. Journal of Neuroscience, 30(47), 15747–15759. https://doi.org/10.1523/JNEUROSCI.2894-10.2010

Ding, L., & Gold, J. I. (2012). Neural correlates of perceptual decision making before, during, and after decision commitment in monkey frontal eye field. Cerebral cortex, 22(5), 1052–1067. https://doi.org/10.1093/cercor/bhr178

Ditterich, J., Mazurek, M. E., & Shadlen, M. N. (2003). Microstimulation of visual cortex affects the speed of perceptual decisions. Nature neuroscience, 6(8), 891–898.

Gao, R., van den Brink, R. L., Pfeffer, T., & Voytek, B. (2020). Neuronal timescales are functionally dynamic and shaped by cortical microarchitecture. Elife, 9, e61277. https://doi.org/10.7554/eLife.61277

Gold, J. I., & Shadlen, M. N. (2007). The neural basis of decision making. Annual review of neuroscience, 30(1), 535–574. https://doi.org/10.1146/annurev.neuro.29.051605.113038

Hanks, T. D., Kopec, C. D., Brunton, B. W., Duan, C. A., Erlich, J. C., & Brody, C. D. (2015). Distinct relationships of parietal and prefrontal cortices to evidence accumulation. Nature, 520(7546), 220–223. https://doi.org/10.1038/nature14066

Hashemi, A., Golzar, A., Smith, J. E. T., & Cook, E. P. (2018). The Magnitude, But Not the Sign, of MT Single-Trial Spike-Time Correlations Predicts Motion Detection Performance. The Journal of Neuroscience, 38(18), 4399–4417. https://doi.org/10.1523/jneurosci.1182-17.2018

Honey, C. J., Thesen, T., Donner, T. H., Silbert, L. J., Carlson, C. E., Devinsky, O., Doyle, W. K., Rubin, N., Heeger, D. J., & Hasson, U. (2012). Slow cortical dynamics and the accumulation of information over long timescales. Neuron, 76(2), 423–434. https://doi.org/10.1016/j.neuron.2012.08.011

Horwitz, G. D., & Newsome, W. T. (1999). Separate signals for target selection and movement specification in the superior colliculus. Science, 284(5417), 1158–1161. https://doi.org/10.1126/science.284.5417.1158

Hunt, L. T., Kolling, N., Soltani, A., Woolrich, M. W., Rushworth, M. F. S., & Behrens, T. E. J. (2012). Mechanisms underlying cortical activity during value-guided choice. Nature neuroscience, 15(3), 470–476. https://doi.org/10.1038/nn.3017

Imani, E., Hashemi, A., Radkani, S., Egger, S. W., & Moazami Goudarzi, M. (2023). The predictive power of intrinsic timescale during the perceptual decision-making process across the mouse brain. bioRxiv, 2023.2001. 2001.522410.

Kim, J.-N., & Shadlen, M. N. (1999). Neural correlates of a decision in the dorsolateral prefrontal cortex of the macaque. Nature neuroscience, 2(2), 176–185. https://doi.org/10.1038/5739

Kobak, D., Brendel, W., Constantinidis, C., Feierstein, C. E., Kepecs, A., Mainen, Z. F., Qi, X.-L., Romo, R., Uchida, N., & Machens, C. K. (2016). Demixed principal component analysis of neural population data. Elife, 5, e10989. https://doi.org/10.7554/eLife.10989

Latimer, K. W., Yates, J. L., Meister, M. L., Huk, A. C., & Pillow, J. W. (2015). Single-trial spike trains in parietal cortex reveal discrete steps during decision-making. Science, 349(6244), 184–187. https://doi.org/10.1126/science.aaa4056

Machens, C. K., Romo, R., & Brody, C. D. (2005). Flexible control of mutual inhibition: a neural model of two-interval discrimination. Science, 307(5712), 1121–1124.

Mante, V., Sussillo, D., Shenoy, K. V., & Newsome, W. T. (2013). Context-dependent computation by recurrent dynamics in prefrontal cortex. Nature, 503(7474), 78–84. https://doi.org/10.1038/nature12742

Mazurek, M. E., Roitman, J. D., Ditterich, J., & Shadlen, M. N. (2003). A role for neural integrators in perceptual decision making. Cerebral cortex, 13(11), 1257–1269. https://doi.org/10.1093/cercor/bhg097

Murray, J. D., Bernacchia, A., Freedman, D. J., Romo, R., Wallis, J. D., Cai, X., Padoa-Schioppa, C., Pasternak, T., Seo, H., & Lee, D. (2014). A hierarchy of intrinsic timescales across primate cortex. Nature neuroscience, 17(12), 1661–1663. https://doi.org/10.1038/nn.3862

Palmeri, T. J., Schall, J. D., & Logan, G. D. (2015). Neurocognitive modeling of perceptual decision making. The Oxford Handbook of Computational and Mathematical Psychology, 320.

Pinto, L., Tank, D. W., & Brody, C. D. (2022). Multiple timescales of sensory-evidence accumulation across the dorsal cortex. Elife, 11, e70263. https://doi.org/10.7554/eLife.70263

Purcell, B. A., Heitz, R. P., Cohen, J. Y., Schall, J. D., Logan, G. D., & Palmeri, T. J. (2010). Neurally constrained modeling of perceptual decision making. Psychological review, 117(4), 1113.

Ratcliff, R. (1978). A theory of memory retrieval. Psychological review, 85(2), 59.

Ratcliff, R., Hasegawa, Y. T., Hasegawa, R. P., Smith, P. L., & Segraves, M. A. (2007). Dual diffusion model for single-cell recording data from the superior colliculus in a brightness-discrimination task. Journal of neurophysiology, 97(2), 1756–1774. https://doi.org/10.1152/jn.00393.2006

Ratcliff, R., & McKoon, G. (2008). The diffusion decision model: theory and data for two-choice decision tasks. Neural Computation, 20(4), 873–922. https://doi.org/10.1162/neco.2008.12-06-420

Ratcliff, R., Smith, P. L., Brown, S. D., & McKoon, G. (2016). Diffusion decision model: Current issues and history. Trends in cognitive sciences, 20(4), 260–281. https://doi.org/10.1016/j.tics.2016.01.007

Roitman, J. D., & Shadlen, M. N. (2002). Response of neurons in the lateral intraparietal area during a combined visual discrimination reaction time task. Journal of Neuroscience, 22(21), 9475–9489. https://doi.org/10.1523/JNEUROSCI.22-21-09475.2002

Rossi-Pool, R., Zainos, A., Alvarez, M., Parra, S., Zizumbo, J., & Romo, R. (2021). Invariant timescale hierarchy across the cortical somatosensory network. Proceedings of the National Academy of Sciences, 118(3). https://doi.org/10.1073/pnas.2021843118

Scott, B. B., Constantinople, C. M., Akrami, A., Hanks, T. D., Brody, C. D., & Tank, D. W. (2017). Fronto-parietal cortical circuits encode accumulated evidence with a diversity of timescales. Neuron, 95(2), 385–398. e385. https://doi.org/10.1016/j.neuron.2017.06.013

Shadlen, M. N., & Newsome, W. T. (2001). Neural basis of a perceptual decision in the parietal cortex (area LIP) of the rhesus monkey. Journal of neurophysiology, 86(4), 1916–1936. https://doi.org/10.1152/jn.2001.86.4.1916

Steinmetz, N. A., Zatka-Haas, P., Carandini, M., & Harris, K. D. (2019). Distributed coding of choice, action and engagement across the mouse brain. Nature, 576(7786), 266–273. https://doi.org/10.1038/s41586-019-1787-x

Usher, M., & McClelland, J. L. (2001). The time course of perceptual choice: the leaky, competing accumulator model. Psychological review, 108(3), 550.

Wang, Q., Ding, S.-L., Li, Y., Royall, J., Feng, D., Lesnar, P., Graddis, N., Naeemi, M., Facer, B., Ho, A., Dolbeare, T., Blanchard, B., Dee, N., Wakeman, W., Hirokawa, K. E., Szafer, A., Sunkin, S. M., Oh, S. W., Bernard, A., . . . Ng, L. (2020). The Allen Mouse Brain Common Coordinate Framework: A 3D Reference Atlas. Cell, 181(4), 936–953.e920. https://doi.org/10.1016/j.cell.2020.04.007

Wang, X.-J. (2002). Probabilistic decision making by slow reverberation in cortical circuits. Neuron, 36(5), 955–968. https://doi.org/10.1016/s0896-6273(02)01092-9

Wong, K.-F., Huk, A. C., Shadlen, M. N., & Wang, X.-J. (2007). Neural circuit dynamics underlying accumulation of time-varying evidence during perceptual decision making. Frontiers in computational neuroscience, 1, 6. https://doi.org/10.3389/neuro.10.006.2007

Wong, K.-F., & Wang, X.-J. (2006). A recurrent network mechanism of time integration in perceptual decisions. Journal of Neuroscience, 26(4), 1314–1328.

Yartsev, M. M., Hanks, T. D., Yoon, A. M., & Brody, C. D. (2018). Causal contribution and dynamical encoding in the striatum during evidence accumulation. Elife, 7, e34929. https://doi.org/10.7554/eLife.34929

Zatka-Haas, P., Steinmetz, N. A., Carandini, M., & Harris, K. D. (2021). Sensory coding and the causal impact of mouse cortex in a visual decision. Elife, 10, e63163. https://doi.org/10.7554/eLife.63163

Zoltowski, D. M., Latimer, K. W., Yates, J. L., Huk, A. C., & Pillow, J. W. (2019). Discrete Stepping and Nonlinear Ramping Dynamics Underlie Spiking Responses of LIP Neurons during Decision-Making. Neuron, 102(6), 1249–1258.e1210. https://doi.org/10.1016/j.neuron.2019.04.031

Zoltowski, D. M., Pillow, J. W., & Linderman, S. W. (2020). Unifying and generalizing models of neural dynamics during decision-making. arXiv preprint arXiv:2001.04571. https://doi.org/10.48550/arXiv.2001.04571

